# Decoding Selective Attention to Context Memory: An Aging Study

**DOI:** 10.1101/324384

**Authors:** Patrick S. Powell, Jonathan Strunk, Taylor James, Sean M. Polyn, Audrey Duarte

## Abstract

Emerging evidence has suggested that the tendency for older adults to bind too much contextual information during encoding (i.e., hyper-binding) may contribute to poorer memory for relevant contextual information during retrieval. While these findings are consistent with theories of age-related declines in selective attention and inhibitory control, the degree to which older adults are able to selectively attend to relevant contextual information during encoding is unknown. To better understand the neural dynamics associated with selective attention during encoding, the current study applied multivariate pattern analyses (MVPA) to oscillatory EEG in order to track moment-to-moment shifts of attention between relevant and irrelevant contextual information during encoding. Young and older adults studied pictures of objects in the presence of two contextual features: a color and a scene, and their attention was directed to the object’s relationship with one of those contexts (i.e., target context). Results showed that patterns of oscillatory power successfully predicted whether selective attention was directed to a scene or color, across age groups. Individual differences in overall classification performance were associated with individual differences in target context memory accuracy during retrieval. However, changes in classification performance within a trial, suggestive of fluctuations in selective attention, predicted individual differences in hyper-binding. To the best of our knowledge, this is the first study to use MPVA techniques to decode attention during episodic encoding and the impact of attentional shifts toward distracting information on age-related context memory impairments and hyper-binding. These results are consistent with the as-of-yet unsubstantiated theory that age-related declines in context memory may be attributable to poorer selective attention and/or greater inhibitory deficits in older adults.

## 1. Introduction

Episodic memories contain information not only about what happened during an event, but also information about contextual details, such as where and when the event happened. These contextual associations are what give memories their episodic quality and allow us to distinguish one event from another. Numerous studies provide converging evidence that episodic memory declines with increasing age (Buckner, 2004; Craik, 1994; Craik and Byrd, 1982; Craik and McDowd, 1987; Dumas and Hartman, 2008; Hess and Blanchard-Fields, 1996), however this decline tends to disproportionately impact memory for contextual details relative to memory of individual items or item memory (Mitchell and Johnson, 2009; Spencer and Raz, 1995). Memory for contextual details is thought to be more reliant on frontally-mediated cognitive control processes compared to item memory, thus greater declines in context memory may in part be related to greater age-related disruption of frontal brain regions. As a result, memory tasks placing high demands on cognitive control (e.g., context memory) are more likely to reveal age-related impairments (Cohn et al., 2008; Duarte et al., 2008).

Although context memory is disproportionately impacted by age, emerging evidence suggests that performance can be improved in both younger and older adults when attention is directed towards task-relevant associations during encoding (Dulas and Duarte, 2013, 2014; Glisky and Kong, 2008; Glisky et al., 2001; Hashtroudi et al., 1994; James et al., 2016; Naveh-Benjamin et al., 2007). That is, when an individual’s attention is intentionally directed toward a relevant item-context association during encoding (i.e., “Is this item likely to be found in this scene?”), context memory performance improves relative to the when attention is solely directed to a single item (i.e., “Is this item larger than a shoebox?”) (Dulas and Duarte, 2013; Glisky and Kong, 2008; Glisky et al., 2001; Hashtroudi et al., 1994; Kuo and Van Petten, 2006; Naveh-Benjamin et al., 2007). Thus, directing attention to the relevant item-context association during encoding may strengthen the relationship between the item and its context and increase the likelihood of successful retrieval.

In everyday situations, we often have multiple features competing for our attention, and our ability to encode some may depend on our ability to successfully ignore others. As aging is well-known to increase susceptibility to interference (Hasher and Zacks, 1988), it is conceivable that context memory impairments may be particularly evident in the presence of task-irrelevant features.

Consistent with this hypothesis, we previously found that older adults were less successful than young adults in selectively encoding relevant item-context associations when distracting context features were present (James et al., 2016; Strunk et al., 2017). One explanation for the reduced benefit in older adults may be a reduction in inhibitory control or the ability to selectively attend to relevant contextual features and ignore irrelevant ones (Campbell et al., 2010; Hasher and Zacks, 1988). During encoding, reduced selective attention in older adults may lead to the formation of overly broad associations such that item-context associations include both relevant and irrelevant contextual features - a process known as *hyper-binding*. Greater hyper-binding in older adults is thought to result in impoverished memory representations for relevant contextual features and increase the conditional dependence between relevant and irrelevant features during retrieval (Boywitt et al., 2012; Meiser et al., 2008; Peterson and Naveh-Benjamin, 2016; Starns and Hicks, 2008). In other words, despite poorer memory performance for relevant contextual features, older adults may be more likely to recover irrelevant contextual features compared to younger adults. Consequently, hyper-binding is more likely to reduce, rather than enhance, older adults’ performance in most explicit memory tasks.

In our previous work involving electroencephalography (EEG) recordings during context retrieval (James et al., 2016; Strunk et al., 2017) we examined the role of selective attention on context memory performance in younger and older adults by explicitly directing attention to the relationship between one of two simultaneously presented contexts (i.e., a color and a scene) during encoding.

Participants in this study were asked to direct attention to the appropriate context (i.e., attended/target context) and ignore the other (i.e., unattended/distractor context). At retrieval, we assessed context memory performance for both the attended and unattended contexts. Behavioral results from this study indicated that both younger and older adults demonstrated better memory for the attended context relative to the unattended context, suggesting both groups were able to selectively attend to the appropriate context during encoding. Yet, older adults showed reduced memory for the attended context and greater conditional dependence between the two contextual features (i.e., hyper-binding) compared to younger adults. Event related potentials (ERPs) indicated that older adults showed a more pronounced late posterior negativity (LPN) than young adults (James et al., 2016). Furthermore, oscillatory EEG results revealed an age-related reduction in early theta synchronization and greater reliance on a late-onset sustained posterior beta desynchronization for successful context memory retrieval (Strunk et al., 2017).

The LPN has been associated with episodic reconstruction through reactivation of context-specifying information (Cycowicz et al., 2001; Johansson and Mecklinger, 2003). Theta synchronization and posterior maximal beta desynchronization have been linked to recollection (Addante et al., 2011; Khader and Rosler, 2011) and sensory reactivation of sought-after perceptual features (for review: Klimesch, 2012; Waldhauser et al., 2016; Waldhauser et al., 2012), respectively. Taken together, these findings suggest a potential consequence of hyper-binding in older adults is greater dependence on episodic reconstruction processes in order to recover relevant contextual information. This fits with the notion that, during encoding, older adults form weaker item-context associations for the attended context. However, the degree to which these weaker associations are the result of poorer selective attention to the attended/target context and/or poorer suppression of unattended/distractor context *during encoding* remains unclear.

To better understand how attentional mechanisms employed by younger and older adults during encoding affects subsequent retrieval of relevant contextual information we examined neural oscillatory signals at encoding in the current study. Using multivariate pattern analysis (MVPA) we assessed category-specific patterns of oscillations during an encoding period when participants were directed to attend to a centrally-presented gray scale object and one of two simultaneously presented contextual features (i.e., color or scene). To determine whether poorer context memory performance in older adults is a consequence of poorer selective attention to the target context and/or poorer suppression of the distractor context, we used MVPA to characterize patterns of neural oscillatory signals that corresponded to a particular context (i.e., scene vs. color) and spatial position (i.e., positioned to the left or right of a centrally-presented object). MVPA differs from univariate analyses in that multiple frequencies and time bins (EEG) are jointly analyzed for their sensitivity to experimental manipulations or memory states (reviewed in Norman et al., 2006).

Prior studies using MVPA techniques with oscillatory EEG data have indicated that category-specific oscillatory patterns during encoding contain information about recently presented items while encoding new ones (Morton et al., 2013; Morton and Polyn, 2017) as well as which category is in the focus of attention (LaRocque et al., 2013; Pereira et al., 2009; Polyn et al., 2005). Category-specific oscillatory patterns have also proven to be reliable predictors of subsequent memory accuracy (Kuhl et al., 2012; Morton et al., 2013; Morton and Polyn, 2017). Category-specific neural representations may occur covertly, such that a data-driven method like MVPA is well-suited to detect the extent of one’s attentional focus towards a particular category in the absence of an overt behavioral response (e.g., controlled eye movements). Hence, MVPA can inform our understanding of the neural mechanisms underlying selective attention to relevant contextual information during encoding. In turn, this may offer additional insight as to whether individual and/or age-related differences in selective attention predict context memory accuracy and hyper-binding as measured during retrieval.

## 2. Methods

### 2.1. Participants

The current study included the participants from two previously published EEG studies (James et al., 2016; Strunk et al., 2017). This included 22 young (18 to 35) and 21 older (60 to 80) healthy, right-handed adults. All participants were native English speakers and had normal or corrected vision. Participants were compensated with course credit or $10 per hour, and were recruited from the Georgia Institute of Technology and surrounding community. No participants reported neurological or psychiatric disorders, vascular disease, or use of any medications affecting the central nervous system. Participants completed a standardized neurological battery and were excluded if their scores fell above or below two standard deviations of the group mean (see **Table 1**). All participants signed consent forms approved by the Georgia Institute of Technology Institutional Review Board. Three older participants were excluded in this analysis because EEG recordings from one or more of the encoding blocks were not available due to computer malfunction.

**Table 1.** Participant Demographics

| *Measure* | Young (n=22) | Old (n=18) |
| --- | --- | --- |
| Age | 21.33 (19.41, 23.25) | 67.86 (66.06, 69.66) |
| Gender (F/M) | 9/13 | 12/6 |
| Education | 14.21 (13.51, 14.91) | 15.23 (14.22, 16.20) |
| Letter Fluency | 46.39 (40.30, 52.48) | 50.82 (40.93, 60.29) |
| List Recall (Immediate) | 10.28 (9.43, 11.13) | 9.17 (7.75, 10.58) |
| List Recall (Delayed) | 11.28 (10.64, 11.91) | 10.11 (8.45, 11.66) |
| Trails A (in seconds) | 23.89 (20.62, 27.16) | 37.22 (25.94, 47.03)** |
| Trails B (in seconds) | 47.45 (41.08, 53.83) | 85.03 (67.30, 102.31)** |
| MoCA (older adults only) | ----- | 27.06 (26.02, 28.10) |

Note: The 95% confidence interval for the mean is in parentheses. Average test scores reported above are mean raw scores

### 2.2. Materials

Four hundred thirty-two grayscale images of objects were collected from the Hemera Technologies Photo-Object DVDs and Google images. During encoding, 288 of these objects were studied, half used when attention was directed to a color and half when directed to a scene. Each grayscale object was centrally presented on the screen on a white background. Positioned to the left and right of the object was a color square and scene. The positions of the context features (e.g., color or scene) were counter-balanced across blocks so that they were presented an equal number of times the on the right and left-hand side of the center object. For each encoding trial, participants were instructed to direct their attention to either the colored square or the scene, which served as the target context for that trial. The possible scenes included a studio apartment, cityscape, or island. The possible colored squares included green, brown, or red. Each of the 42 context and object images spanned a maximum vertical and horizontal visual angle of approximately 3˚. During retrieval, all 288 objects were included in the memory test along with 144 new object images that were not studied during encoding. Study and test items were counterbalanced across subjects.

### 2.3. Procedure

**Figure 1** illustrates the procedure used during the study and test phases. Prior to each phase, participants were provided instructions and given 10 practice trials. For the study phase, participants were asked to make a subjective yes/no judgment about the relationship between the object and either the colored square (i.e., is this color likely for this object?) or the scene (i.e., is this object likely to appear in this scene?). Instructions for the task stated that on any particular trial the participant should attend to one context and ignore the other context. Within the study phase there were four blocks; each block was divided into four mini-blocks containing 18 trials each (see **Figure 2**). Prior to beginning each mini-block, participants were provided a prompt (e.g., “Now you will judge how likely the color is for the object” or “Now you will judge how likely the scene is for the object”). Because prior evidence has suggested that memory performance in older adults is more disrupted when having to switch between two different types of tasks (Kray and Lindenberger, 2000), mini-blocks were included to not only orient the participant to which context they should pay attention to in the following trials, but also to reduce the task demands of having to switch from judging one context (e.g., color) to judging the other (e.g., scene). Additionally, each trial within a mini-block included a reminder prompt presented underneath the images during study trials (see **Figure 1**).

**Figure 1.**
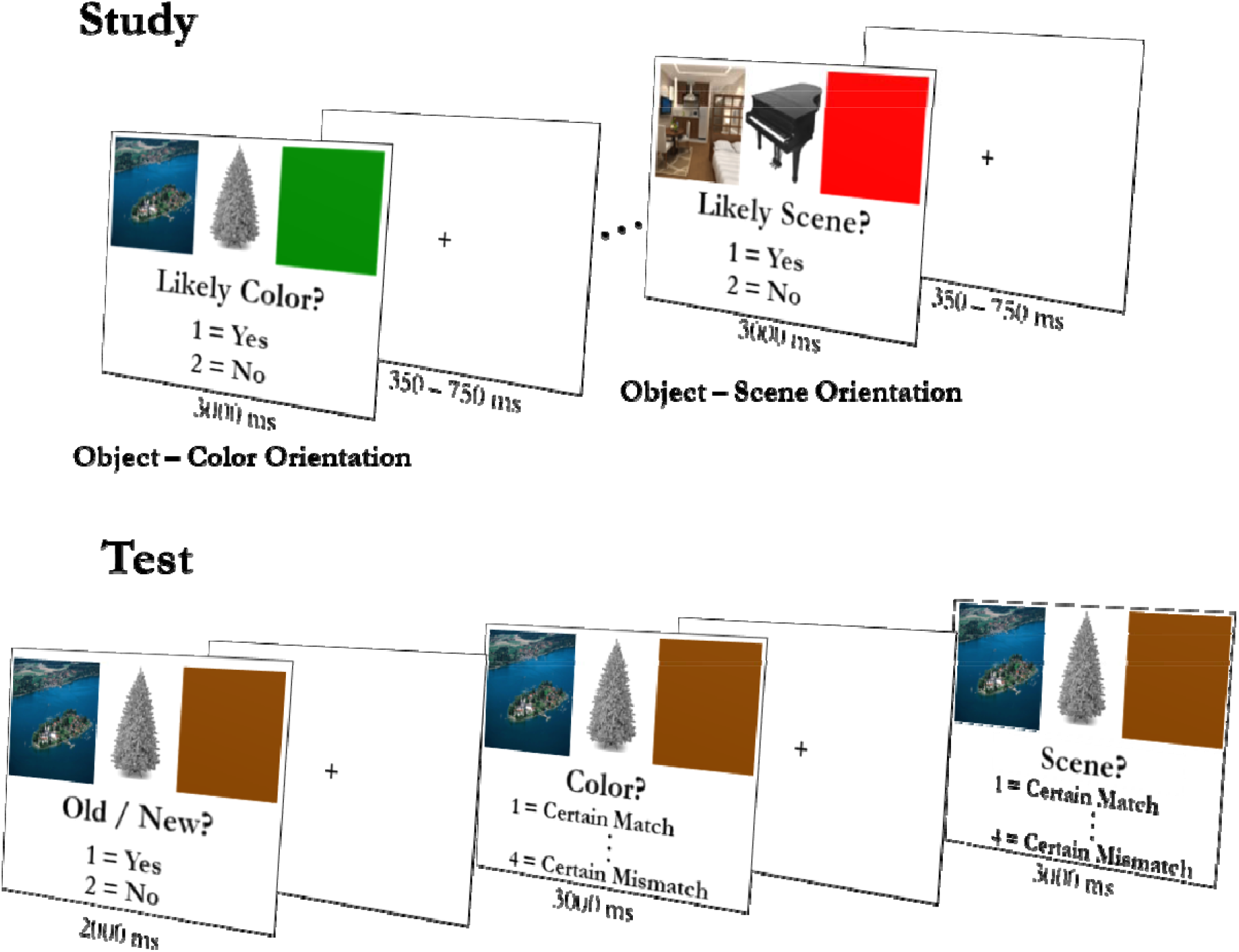
Task design for study and test phase

**Figure 2.**
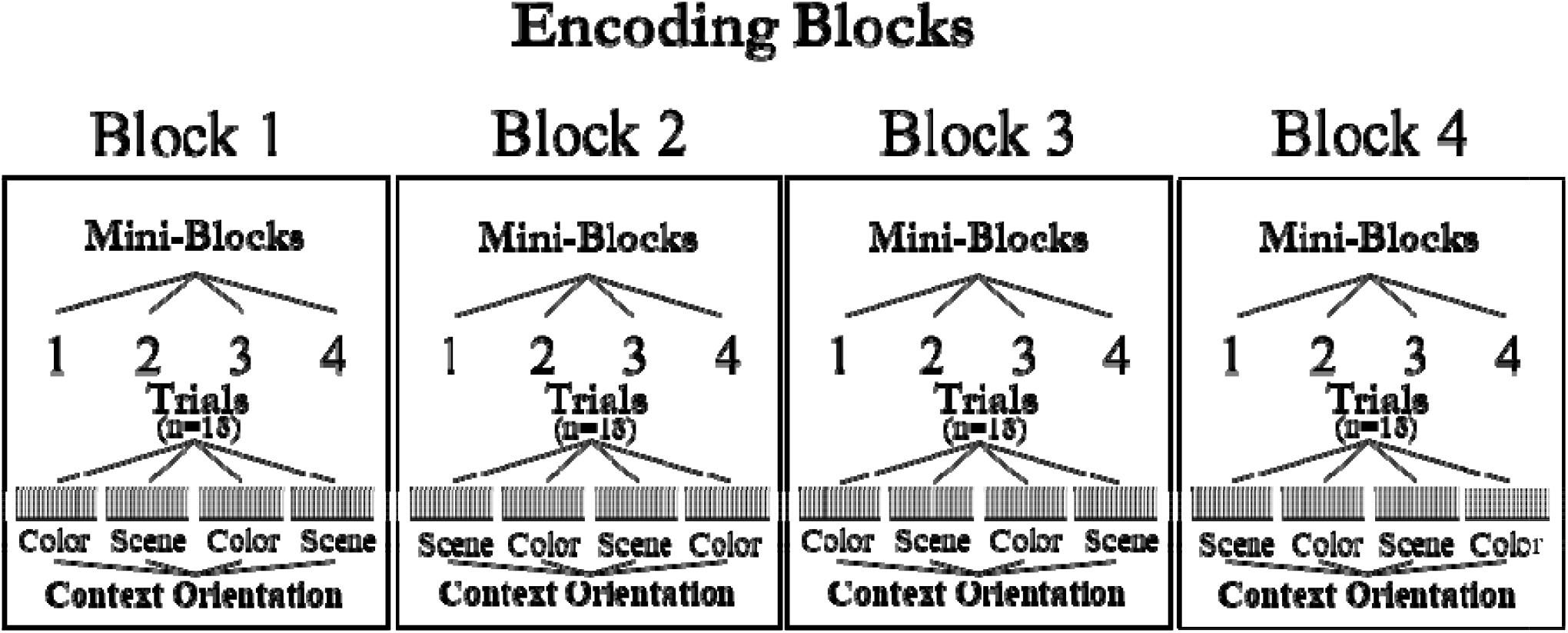
Mini-blocks used in cross-validation (n-1). Four mini-blocks per block, 18 trials per mini-block.

During test, participants were presented with both old and new objects. As in the study phase, each object was flanked by both a scene and a colored square. For each object, the participant first decided whether it was an old or a new image. If the participant responded that it was new, the next trial began after 2000 ms. If participants responded that it was old, then they were asked to make two additional judgments about each context feature (i.e., one about the colored square and another about the scene). The order of these last two questions was counterbalanced across participants. For old items, the pairing was arranged so that an equal number of old objects were presented with: (1) both context images matching those presented at encoding, (2) only the scene matching, (3) only the color matching, and (4) neither context images matching. Responses to the context questions were made on a scale from 1 (certain match) to 4 (certain mismatch). For those items correctly identified as ‘Old’, we classified a ‘Target Context-Hit’ as correctly identifying whether the attended context (scene or color) was the same as or different from encoding, regardless of memory for the unattended context. An incorrect response was classified as a ‘Target Context-Miss’. In total, there were four study and four test blocks. Young adults completed all four study blocks before the four test blocks. For older adults, in order to better equate item memory performance with young adults, the memory load was halved so that they completed a two-block study-test cycle twice (two study, two test, two study, two test). Both younger and older adults completed a short practice of both the study and test blocks before starting the first study block. Thus, both younger and older adults knew of the upcoming memory test.

### 2.3. EEG recording

Continuous scalp-recorded EEG data were collected from 32 Ag-AgCl electrodes using an ActiveTwo amplifier system (BioSemi, Amsterdam, Netherlands). Electrode position follows the extended 10-20 system (Nuwer et al., 1998). Electrode positions included: AF3, AF4, FC1, FC2, FC5, FC6, FP1, FP2, F7, F3, Fz, F4, F8, C3, Cz, C4, CP1, CP2, CP5, CP6, P7, PO3, PO4, P3, Pz, P4, P8, T7, T8, O1, Oz, and O2. External left and right mastoid electrodes were used for referencing offline. Two electrodes placed superior and inferior to the right eye recorded vertical electrooculogram (VEOG) and two additional electrodes recorded horizontal electrooculogram (HEOG) at the lateral canthi of the left and right eyes. EEG was sampled at 1024 Hz with 24-bit resolution without high or low pass filtering.

### 2.4. EEG preprocessing

Offline analysis of the EEG data was done in MATLAB 2015b with the EEGLAB (Delorme and Makeig, 2004), ERPLAB (Lopez-Calderon and Luck, 2014), and FIELDTRIP (Oostenveld et al., 2011) toolboxes. The continuous data were down sampled to 256 Hz, referenced to the average of the left and right mastoid electrodes, and band pass filtered between .5 Hz and 125 Hz. The data were then epoched from −1000 ms prior to stimulus onset to 3000 ms. The time range of interest was set to −300 ms to 2000 ms, but a longer epoch is needed to account for signal loss at both ends of the epoch during wavelet transformation. Each epoch was baseline corrected to the average of the whole epoch, and an automatic rejection process removed epochs with extreme voltage shifts that spanned across two or more electrodes, or epochs in which a blink occurred during stimulus onset. The automated rejection processes identified epochs with the following parameters in the raw data: 1) The voltage range was greater than 99th percentile of all epoch voltage ranges within a 400 ms window (sliding in 100 ms intervals across each epoch). 2) The linear trend slope exceeded the 95th percentile of all epoch ranges with a min R^2^ value of 0.3. 3) The voltage range was greater than 95th percentile of all epoch voltage ranges within a 100 ms window (sliding in 25 ms intervals across each epoch), between −150 and 150 ms from stimulus onset for frontal and eye electrodes only. Then an independent component analysis (ICA) was run on all head electrodes in order to identify additional artifacts highlighted by the components. The following parameters were used on the components to reject epochs: 1) The voltage range was greater than 99th percentile of all epoch voltage ranges within a 400 ms window (sliding in 100 ms intervals across each epoch). 2) The kurtosis or joint probability exceeded 15 standard deviations within the component or 23 standard deviations of all components for the epoch. In order to identify activity related to ocular artifacts (i.e., blinks and horizontal eye movements), ICA was run on the first 20 principle components of the head electrodes for the accepted epochs. Components related to ocular artifacts were removed from the data by visually inspecting the topographic component maps and component time course with the ocular electrodes (Bell and Sejnowski, 1995; Delorme et al., 2007; Hoffmann and Falkenstein, 2008). Since the epochs were no longer baselined to a specific time period after removing components related to ocular activity, each epoch was re-baselined to the −300 to −100 ms time period before stimulus onset. This was solely for the purposes of visual inspection and detection of additional artifacts in each epoch (e.g., amplifier saturation, spiking, extreme values, uncorrected ocular activity), and does not affect the frequency decomposition. If a dataset contained a noisy electrode (e.g., greater than 30% of the data needed to be rejected), it was removed from the processing stream and interpolated using the surrounding channels to estimate the activity within the bad channel before running the time frequency process (Delorme and Makeig, 2004). After all processing steps, approximately 13% (SD = 8%) of the epochs were removed.

### 2.5. Frequency decomposition

Each epoch was transformed into a time frequency representation using Morlet wavelets (Percival and Walden, 1993) with 78 linearly spaced frequencies from 3 and 80 Hz, at 5 cycles. During the wavelet transformation, each epoch was reduced to the time range of interest and down sampled to 50.25 Hz (Cohen, 2014). For the following MVPA analyses, we examined item hit events across both context features (i.e., attend color and attend scene), including all levels of confidence. The average number of trials for younger and older adults are as follows: Younger (*M* = 190.50, *SD* = 41.01); Older (*M* = 177.06, *SD* = 38.56).

### 2.5 Multivariate pattern analysis

We used multivariate pattern analysis (Norman et al., 2006) to decode stimulus category based on patterns of oscillatory power at encoding (Morton et al., 2013; Newman and Norman, 2010). Classification was carried out using penalized logistic regression (penalty parameter = 10), using L2 regularization (Duda et al., 2001). Classification analyses were conducted using Aperture (available at: http://mortonne.github.io/aperture/) and the Princeton MVPA Toolbox (available at: http://www.pni.princeton.edu/mvpa). Using the distributed pattern of oscillatory power during each epoch, a cross-validation procedure was used to train the classifier to discriminate conditions by target context feature (color or scene) and spatial location (target context positioned to the left or right of the item). The classifier was trained on all but one mini-block, then applied to the epochs on the left-out list to evaluate its performance, measured as the fraction of items classified correctly (**Figure 2**).

Performance at each mini-block was then averaged to produce a measure of overall prediction accuracy. This procedure was repeated separately for each participant.

Several sets of study-phase patterns were created for the analyses reported below. For our first analysis, we wanted to simply characterize the particular frequencies and time windows containing robust category-specific activity. In order to do so, we examined all 78 frequencies (3-80 Hz) and 46 time bins (~45 ms per bin) **(Figure. 4**). The values for each feature reflect oscillatory power at each time-frequency pairing and taken from trials during the study period where the participant correctly identified the correct object (item – hit) at retrieval. Average oscillatory power from trials where the participant failed to correctly recognize a previously studied item (i.e., Misses) were excluded because context memory test trials were only presented after a participant identified an object as previously seen before (i.e., Old). Since this initial classification analysis was intended for illustrative purposes only (**Figure 4**), results from this classification were not submitted to additional statistical analyses.

**Figure 3.**
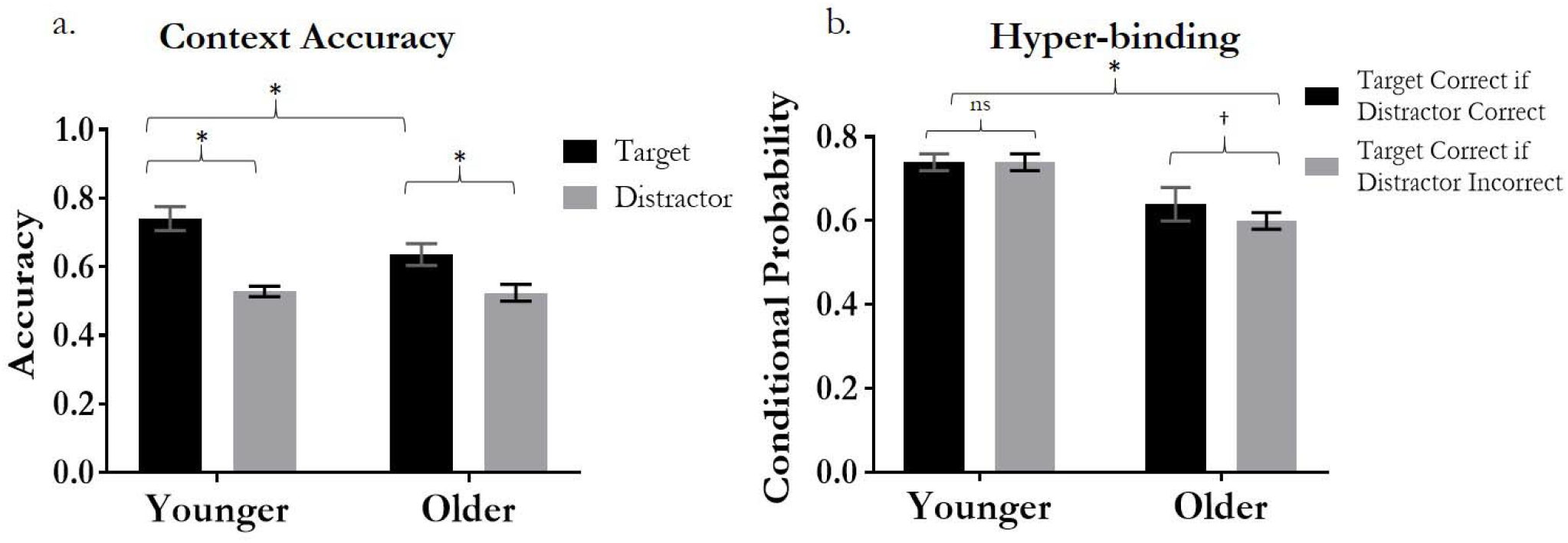
Context memory accuracy and hyper-binding in younger and older adults. * indicates significant difference (p < .01), † indicates a marginally significant difference (p = .10), ns indicates non-significant. Error bars represent 95% confidence interval.

**Figure 4.**
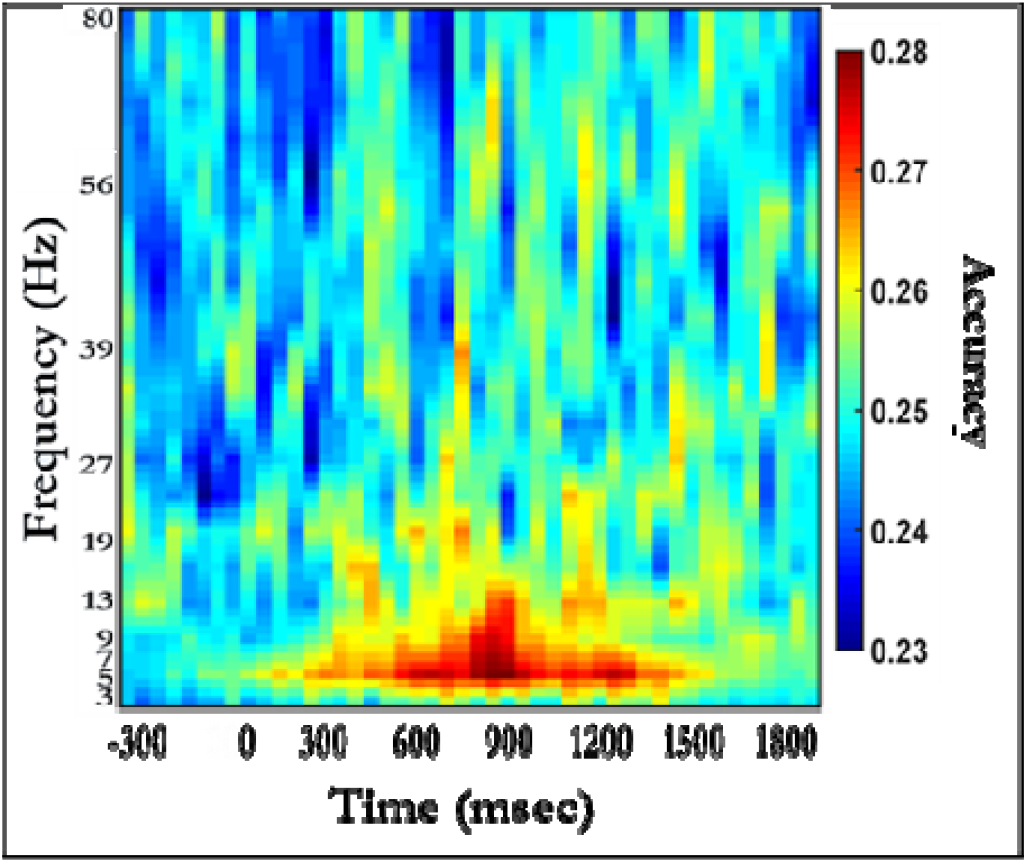
Classifier accuracy for separate cross-validation analyses at 46 time bins and all frequencies during encoding. Stimuli were classified based on oscillatory power over all electrodes. Light blue corresponds to chance performance (0.25).

To more closely examine how classification performance varied as a function of time and frequency, a second across-electrode pattern was created by binning the range of frequencies into the five frequency bands of interest: Delta (3-4Hz) Theta (4-7 Hz), Alpha (8-12 Hz), Beta (14-26 Hz), and Gamma (27-80 Hz); and expanding the time bins to create eight time bins, approximately 300 ms per bin, ranging from −300ms prior to stimulus onset to 2000 ms post-stimulus onset (e.g., −300-0, 0-300, 300-600, 600-900, 900-1200, 1200-1500, 1800-2000 ms). Importantly, these longer time bins and frequency bands were not selected based on the results from the previous classification analysis. Rather, the frequency bands were selected a priori based on standard frequency bands used in prior literature (Bastiaansen et al., 2012; Klimesch, 1999). Likewise, the specific time bins are similar to those used in previous studies (Morton et al., 2013; Morton and Polyn, 2017), and allowed for symmetrical sampling of classification performance over the trial epoch while at the same time reducing the number of comparisons. However, one might wonder if the selection of eight 300ms time bins could potentially rendered more favorable classification results. To address this, we conducted additional classification analyses selecting a larger number of time-bins as features. Results from these analyses indicated that classification performance was not sensitive to selection of specific time windows^1^. As such, the findings reported below were based on pattern classification of average oscillatory power across all electrodes from each of the eight 300-ms bins and five frequency bands. The same cross-validation procedure was used (i.e., training on all but one mini-block) and an across-electrode pattern was generated for each context category, again where the value for each feature of the pattern was oscillatory power taken from each of the 40 time bin-frequency band pairings (e.g., eight time bins and the five frequency bins - Delta, Theta, Alpha, Beta, and Gamma). As previously mentioned above, the trials included in this analysis were trials where the participant correctly identified the correct object (item – hit) at retrieval. Trials where the participant failed to correctly recognize a previously studied item were excluded because context memory was not tested.

Following the analysis of the across-electrode pattern with oscillatory power at eight time bins (-300-0, 0-300, 300-600, 600-900, 900-1200, 1200-1500, 1800-2000 ms) and five frequency bands [Delta (3-4Hz) Theta (4-7 Hz), Alpha (8-12 Hz), Beta (14-26 Hz), and Gamma (27-80 Hz)].used as features, we carried out a permutation test to determine whether classification performance was significantly above chance for each subject. For each subject, the category context labels used in the cross-validation described in the text were scrambled, and the mean classification performance metric was calculated for each subject. This process was repeated 5000 times to establish a null distribution of performance metric scores, and performance was considered significant if the observed score was >95% of the null distribution. As can be seen in the supplementary material (**Figure S1**), mean noise-level classification performance was .25 with a 95% confidence interval of .2496 to .2503. This familywise null distribution was then used to set the significance threshold when determining if classification performance was significantly above chance (Sederberg et al., 2003).

To evaluate the statistical reliability of classification performance from this pattern, we conducted a Time Bin x Frequency Band x Age Group mixed factorial repeated-measures ANOVA. Time bin and frequency band were treated as the within-subject factors, age group as the between-subjects factor, and classification performance as dependent variable. If a significant main effect emerged, we then conducted one-sample *t*-tests against chance (25%) to identify the time bins and frequency bands in which classification performance successfully identified patterns of oscillatory power that differentiated between contextual features and spatial location. All post-hoc analyses were corrected for multiple comparisons using the Holm-Bonferroni correction (Holm, 1979).

A classification analysis was also conducted to determine the category discriminability of oscillatory activity at specific electrodes. This classification identified a number of electrodes with high classification performance (for additional details see supplementary material). Subsequent analysis used average linkage clustering to sort electrodes based into high- and low-performance clusters. Examining the average difference in classification performance between each pair of electrodes, this technique identified a number of high-performing posterior electrodes (CP5, P3, P7, PO3, Pz, O1, Oz, O2, PO4, P8, P4, and CP6). Given the nature of the task, and previous work indicating laterality differences in EEG activity associated with stimulus category and spatial location (van der Lubbe et al., 2000; van der Lubbe and Utzerath, 2013; Wascher and Wauschkuhn, 1996), it is possible that classification performance within these electrodes might vary as a function of whether attention to a particular contextual feature was contralateral or ipsilateral to the posterior electrodes. Therefore, this cluster was divided into left (CP5, P3, P7, PO3, O1) and right (O2, PO4, P8, P4, CP6) posterior regions of interest (ROIs), excluding two electrodes on the midline (Pz, Oz), to create a final classification pattern where the value for each feature of the pattern was average oscillatory power within these two posterior clusters, five frequency bands, and three time bins (0-500, 500 – 1000, 1000 – 2000 ms). Based on findings from our earlier classification analysis using eight 300 ms time bins, we selected slightly larger time bins (~500 ms per bin) to examine within-trial changes in classification performance at early, middle, and late time windows. Again, statistical reliability of this pattern classification was evaluated using a Time Bin x Frequency Band x Hemi-field (contra-vs. ipsilateral to target context) x Age Group mixed factorial repeated-measures ANOVA. Time bin, frequency band, and hemi-field were treated as the within-subject factors, age group as the between-subjects factor, and classification performance as the dependent variable.

Following the evaluation of classification performance from each of the two classification analyses described above, several analyses examined the relation between classification performance and context memory accuracy. A brief description of the statistical approach used in these analyses is provided prior to the reported findings.

## 3. Results

### 3.1. Behavioral Results

Item recognition accuracy was estimated using the Pr measure of discriminability: p(hits) –p(false alarms). Pr estimates for younger and older adults were, *M* = 0.67, *SD* = 0.15; *M* = 0.61 *SD* = 0.15, respectively. No significant group difference in item recognition was found [*t*(41) < 1]. Chance performance for context accuracy was 0.50. For both age groups, context accuracy was above chance for the target contexts (Younger: *M* = .74, *SD* = .08; Older: *M* = .66, *SD* = .07, *ts* > 9.05, *ps* < 0.001). Context accuracy was above chance for the distractor context in the young [*M* = .53, *SD* = .03, *t*(21) = 3.93, *p* = .001], but not the old [*M* = .52, *SD* = .05, *t*(17) = 2.04, *p* = .06]. A Context Feature (target vs. distractor) x Age Group (younger vs. older) ANOVA revealed main effects of context feature, *F*(1, 38) = 131.66, *p* < .0001, ƞ_p_^2^ = .78, and age group, *F*(1, 38) = 19.14, *p* < .001, ƞ_p_^2^ = .34, which were qualified by a significant interaction between these factors, *F*(1, 38) = 12.70, *p* <.001, ƞ_p_^2^ = .25. Closer inspection of the main effect of context feature indicated that participants were more likely to correctly endorse the target context (*M* = .69; *SD* = .09) compared to the distractor context (*M* = .53; *SD* = .04), suggesting manipulation of attention during encoding was effective at enhancing memory accuracy for the target context. However, the significant context feature by age group interaction that suggested context memory accuracy in older adults, relative to younger adults, was particularly impaired for the target context (**Figure 3a**).

Previously reported findings provided evidence that poorer accuracy for the target context in older adults is consistent with poorer selective attention and greater binding of both the target and distractor context to the object during encoding; which was assessed by examining conditional probabilities of correctly endorsing the target and distractor context (see James et al., 2016; Strunk et al., 2017). Specifically, conditional probabilities were computed via the probability of correctly endorsing the target context given the distractor context was correctly endorsed, p(both correct)/[p(both correct) + p(distractor only correct)], and the probability of correctly endorsing the target context given the distractor context was incorrect was computed as p(target correct)/[p(target correct) + p(neither correct)]. Previous studies have used similar calculations to assess conditional context accuracy (Uncapher et al., 2006). To examine these probabilities, we conducted an Age Group (young, old) x Target Context Probability [p(target correct | distractor correct) vs. p(target correct | distractor incorrect)] repeated measures ANOVA. This revealed a significant main effect of age group, *F*(1, 38) = 21.45, *p* < .001, ƞ_p_^2^ = .36, however the main effect for target context probability and the Age Group x Target Context Probability interaction were non-significant [*F*(1, 38) = 2.05, *p* = .16, ƞ_p_ ^2^ = .05; *F*(1, 38) = 1.31, *p* = .26, ƞ_p_^2^ = .03, respectively]. Despite the non-significant Target Context Probability x Age Group interaction, **Figure 3b** shows that young adults’ ability to correctly identify the target context was unaffected by accuracy of the distractor context. However older adults showed a trend of improved memory for the target if the distractor was also correct, relative to when the distractor was incorrect. In our previous work with this sample (James et al., 2016), older adults exhibited a significant, albeit small, hyper-binding effect. Given the modest effect size reported in this previous work, it is likely that the slightly smaller sample of older adults included in this paper (n = 18) is likely to have resulted in the marginal hyper-binding effect reported here.

### 3.2. Classification Performance

Our initial across-electrode pattern used oscillatory activity taken from 78 frequencies (3-80 Hz) and 46 time bins (~45 ms per bin) as features, where the value of each feature was oscillatory power at each time bin-frequency pairing. This allowed us to characterize which frequencies and time relative to stimulus onset contained information about stimulus category. Results from this analysis (see **Figure 4**) indicated greater classification performance approximately 300 to 1200ms after stimulus onset and primarily in lower frequencies (i.e., 2 to 20Hz), suggesting peak performance in the Delta, Theta, and Alpha frequency bands.

To more closely examine how classification performance varied as a function of time and frequency, a second across-electrode pattern was created by binning the range of frequencies into the five frequency bands of and interest (Delta, Theta, Alpha, Beta, and Gamma) and expanding the time bins to approximately 300 ms per bin (e.g., −300-0, 0-300, 300-600, 600-900, 900-1200, 1200-1500, 1500-1800, & 1800-2000 ms). As previously mentioned, the value for each feature of this pattern was oscillatory power taken from each of the 40 time bin-frequency band pairings (e.g., eight time bins and the five frequency bands - Delta, Theta, Alpha, Beta, and Gamma) for trials where the participant correctly identified the correct object (item – hit) at retrieval. A separate cross-validation classification analysis was conducted on this pattern to examine how classification performance varied by time and frequency. Classification performance taken from each time bin and frequency band was then used in a 2 × 8 × 5 (Age Group [young, old] x Time Bin [-300 to 2000ms] x Frequency Band (delta, theta, alpha, beta, gamma]) repeated measures ANOVA with a polynomial contrast to determine whether classification performance varied between age groups, time bins, and frequency bands.

Results from this analysis indicated significant quadratic effects for time, *F*(1,37) = 25.40, *p* <.001, ƞ_p_^2^ = .41, and frequency, *F*(1,37) = 8.68, *p* = .006, ƞ_p_^2^ = .19. There was no main effect of age, *F* (1,37) = 1.84, *p* = .18, ƞ_p_^2^ = .05, suggesting similar classification performance between younger and older adults. There were no significant 2-way or 3-way interactions between time, frequency, and/or age (all *p*s > .22). Since classification performance did not significantly differ between younger and older adults, classification performance was averaged across younger and older adults for post-hoc analyses.

As can be seen in **Figure 5a**, average classification performance prior to and slightly after stimulus onset (roughly −300 to 300ms) was not significantly above chance (25%). However, after 300 ms performance started to increase, peaking approximately 600ms to 1200ms after stimulus onset. After correcting for multiple comparisons, post-hoc analyses of classification performance at each time bin indicated performance that was significantly greater than chance at 600ms (*M* = .28, *SD* = .05) and 900ms (*M* = .27, *SD* = .04). Given that these performance values reflected classification performance averaged across frequency bands, a follow-up inspection of this pattern was conducted to separately examine classification performance for each frequency band within this peak time window (300-1200ms). Importantly, classification performance illustrated in **Figure 5b** does not reflect a separate classification analysis, rather the purpose of this examination was to determine which frequency band, or bands, contributed to this peak in classification performance. This examination showed relatively high performance for a number of frequency bands including Delta, Theta, Alpha, and Beta frequency bands (**Figure 5b**). After correcting for multiple comparisons, post-hoc analyses indicated classification performance in Delta, Theta, Alpha, and Beta bands was significantly greater than chance (*p*s < .05)^2^.

**Figure 5.**
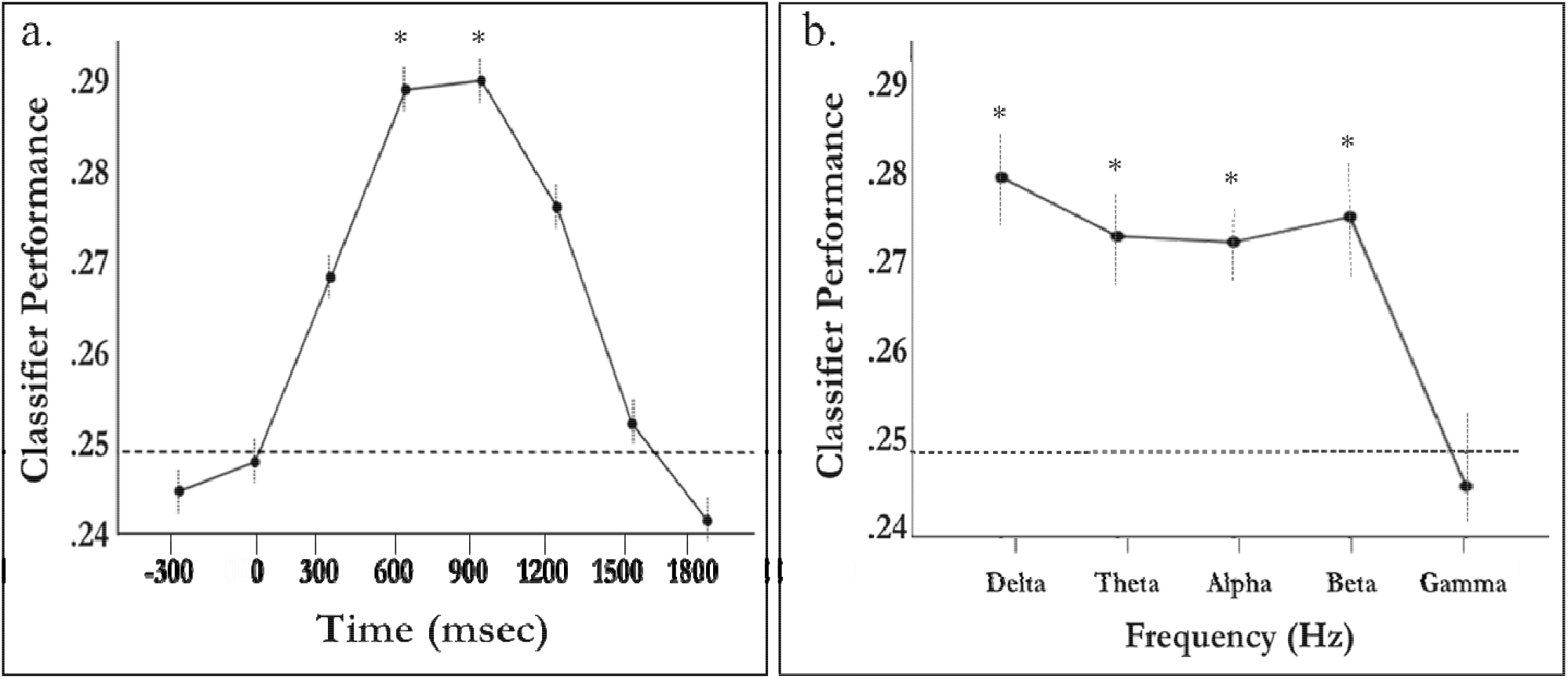
(a) Classifier performance averaged across frequency bands and plotted against time after stimulus onset. Category-specific patterns peak at about 600 to 1200 ms after stimulus onset. (b) Average classifier performance, between 300 and 1200ms, for each frequency band. The dotted line indicates chance performance (0.25) and asterisks denote performance statistically greater than chance after correcting for multiple comparisons (Holm-Bonferroni). Error bars represent within-subject standard error +/-1.5.

### 3.3. Relation between Classification Performance and Context Memory

To determine whether classification performance during encoding reflected selective attention to the association between the item and target context, we first took average classification performance across the trial epoch (0 to 2000 ms) for each frequency band and each participant. These averages were then used to calculate Pearson’s correlation coefficients between average classification performance at each frequency band and a number of behavioral measures of recognition accuracy: (a) *item memory accuracy*-Pr(Hit-False Alarms), (b) *target context accuracy*-the proportion of hits for which participants correctly judged both contexts (i.e., attended and unattended) or only the attended context correctly, (c) *distractor context accuracy*– measured by the proportion of hits for which participants judged only the unattended context correctly, and (d) a composite index of *hyper-binding*^3^.

After correcting for multiple comparisons, results from theses analyses indicated a significant and positive relationship between classification performance in the Beta band and correct recognition of the target context, suggesting greater classification performance during encoding was associated with better target context accuracy (**Table 2**). Similar positive correlations were observed in the Alpha and Gamma bands, but these were marginally significant (*p* = 0.07, *p* = 0.08, respectively). In contrast, significant negative correlations were observed between distractor context accuracy and classification performance in Alpha and Beta bands (after correcting for multiple comparisons), suggesting greater classification performance in these frequency bands was associated with poorer recognition accuracy for the distractor context. Marginally significant negative correlations were observed in the Theta and Gamma bands (*p*s = .06), and there were no significant relationships between classification performance and hyper-binding (*ps* ≥ .25). Importantly, classification performance was not significantly related to item memory (*ps* ≥ .11).

**Table 2.** Correlational coefficients between classification performance at each frequency band and memory accuracy

|  | Delta | Theta | Alpha | Beta | Gamma |
| --- | --- | --- | --- | --- | --- |
| Item Memory | .21 | .26 | .17 | .00 | -.14 |
| Target Context | -.01 | .17 | .29 | .42* | .28 |
| Distractor Context | -.16 | -.40 | -.50* | -.51* | -.35 |
| Hyper-binding | -.18 | .02 | .03 | .01 | -.13 |

Note: \* Significant after correcting for multiple comparisons using Holm-Bonferroni correction
<sup>3</sup> The hyper-binding index was computed by subtracting the conditional probability of correctly endorsing the target context given the distractor was correct, $p(\text{Both correct})/[p(\text{Both correct}) + p(\text{Distractor correct})]$ , by the conditional probability of endorsing the target context given the distractor context was incorrect $p(\text{Target correct})/[p(\text{Target correct}) + p(\text{Neither correct})]$ . Thus, positive values indicate a greater likelihood of correctly recognizing both the target and distractor (i.e., hyper-binding).

Follow-up regression analyses were conducted to determine whether these relationships varied as a function of age. Separate hierarchical linear regressions were conducted on recognition accuracy of the target context and distractor context. For each regression, age group (younger vs. older) was entered as the first categorical predictor. This was followed by classification performance for each frequency band and an Age Group x Classification Performance interaction term, each being entered sequentially to determine their unique impact on context memory accuracy after accounting for other variables in the model. Given the number of comparisons, *p*-values were subjected to the Holm–Bonferroni correction for multiple comparisons to reduce the probability of Type I errors (Holm 1979). Significant results are presented in Table 3. Table S1 in the supplementary material provides standardized regression coefficients for each frequency band.

**Table 3.** Standardized beta coefficients and *p*-values with age and frequency band as predictors of target and distractor context accuracy

| Predictors | Target Context |  | Distractor Context |  |
| --- | --- | --- | --- | --- |
| | $\beta$ | *p* | $\beta$ | *p* |
| Age Group | -.58 | <.001* | .32 | .05 |
| *Classification Performance* |  |  |  |  |
| Alpha | .20 | .15 | -.46* | .002 |
| Beta | .30 | .02 | -.46* | .003 |
| *Age Group by Classification Performance* |  |  |  |  |
| Age Group x Alpha | .20 | .45 | -.53 | .06 |
| Age Group x Beta | .03 | .93 | -.35 | .24 |

Note: \* Significant after correcting for multiple comparisons

As shown in Table 3, results from these regression analyses indicated that after correcting for multiple comparisons and controlling for age, classification performance in the Beta band was a marginally significant positive predictor of target context accuracy, R^2^_change_ = .09, *F*(1,37) = 5.48, β = .30, *p*_corrected_ = .10. Classification performance in the Alpha band was not a significant predictor of target context accuracy after controlling for age, R^2^_change_ = .04, *F*(1,37) = 2.18, β = .20, *p* = .15. In contrast, classification performance in both the Alpha and Beta bands significantly predicted poorer distractor context accuracy, R^2^_change_ = 0.20, *F*(1,37) = 10.75, β = −.46, *p*_corrected_ = R^2^_change_ = 0.20, *F*(1,37) = 10.32, β = −0.46, *p*_corrected_ = .02, respectively. Finally, there were no significant Age Group x Classification Performance interactions (*F*s < .58, *ps* ≥.45).

**Figure 6** further illustrates the relationships between classification performance in the Beta band to item, target, and distractor context accuracy. Specifically, both younger and older adults showed no relationship between classification performance in the Beta band and item memory (**Figure 6a**), and similar positive relationships between classification performance and target context accuracy (**Figure 6b**). Although older adults seemed to exhibit a stronger negative relationship between classification performance and distractor context accuracy (**Figure 6c**), the magnitude of this relationship was not significantly different from that of younger adults (see Table 3). Nevertheless, these findings suggest that during encoding, classification performance in both younger and older adults may reflect better selective attention and encoding of the target context (**Figure 6b**), or, possibly, greater inhibitory control or suppression of attention towards the distractor context (**Figure 6c**).

**Figure 6.**
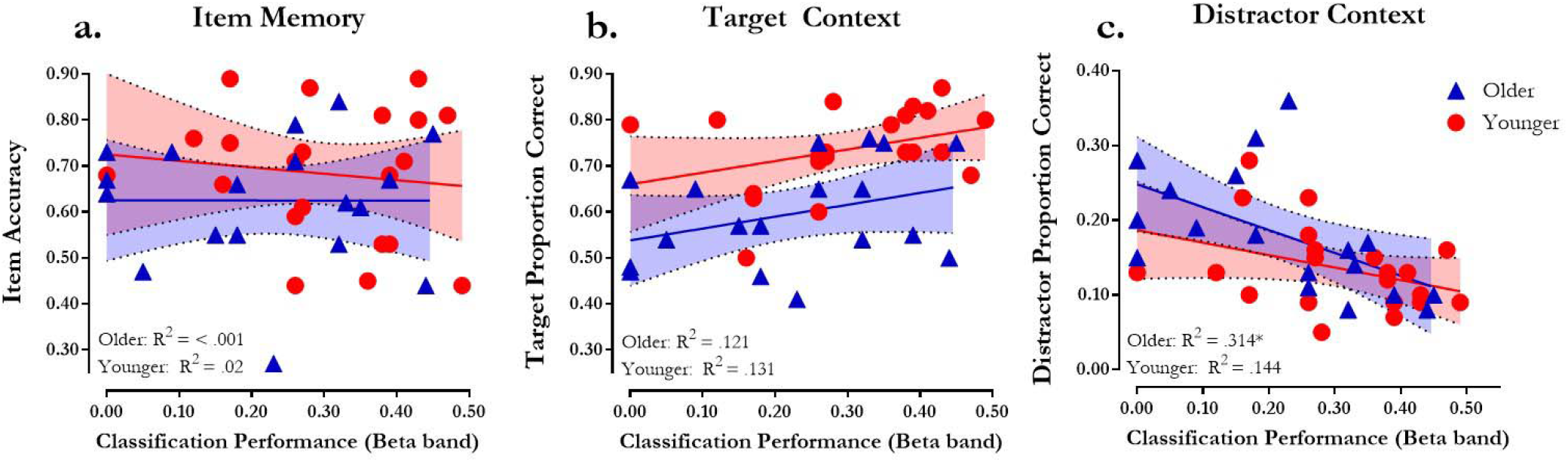
Classifier performance in the beta band with (a) item memory accuracy - measured by p(hits) - p(false alarms), (b) target proportion correct - percentage of trials on which participants judged both a studied item old (hit) and target context accurately, and (c) distractor proportion correct - percentage of trials on which participants judge a studied item old (hit) and only the distractor context accurately. * indicates p-values < .05.

### 3.4. Classification Performance and Spatial Position

We next examined whether patterns of oscillatory power within specific electrode clusters could reliably differentiate when the target context was positioned to the left or right of the centrally presented object (i.e., whether classification performance changed as a function of whether the particular cluster was contra- or ipsilateral to the target context). As previously mentioned, average linkage clustering identified a cluster of high-performing posterior electrodes which were divided into left and right ROIs to examine laterality differences when attention to the target context was directed to the right or left. **Figure 7** shows the electrodes included in left cluster (CP5, P7, PO3, P3, and O1) and the corresponding electrodes in the right cluster (CP6, P8, PO4, P4, and O2). Since these ROIs were selected based on classifier performance for the individual electrodes, it is possible that mean classification performance in these ROIs may be inflated. However, because the ROIs were selected based on data aggregated over all conditions, it is unlikely to bias subsequent analysis of differences between experimental conditions. Therefore, we included the two clusters as features in this classification along with five frequency bands (Delta, Theta, Alpha, Beta, Gamma) and three time bins, designated hereafter as Time 1 (0 to 500 ms), Time 2 (500 to 1000 ms), and Time 3 (1000 to 2000 ms). Similar to previous analyses, the value for each feature of this pattern was oscillatory power taken from each of the 30 cluster-time-frequency pairings (e.g., two clusters, three time bins, and the five frequency bands - Delta, Theta, Alpha, Beta, and Gamma) for trials where the participant correctly identified the correct object (item – hit) at retrieval. These time bins were selected in order to examine classification performance within each electrode cluster at early, middle, and late time windows^4^.

**Figure 7.**
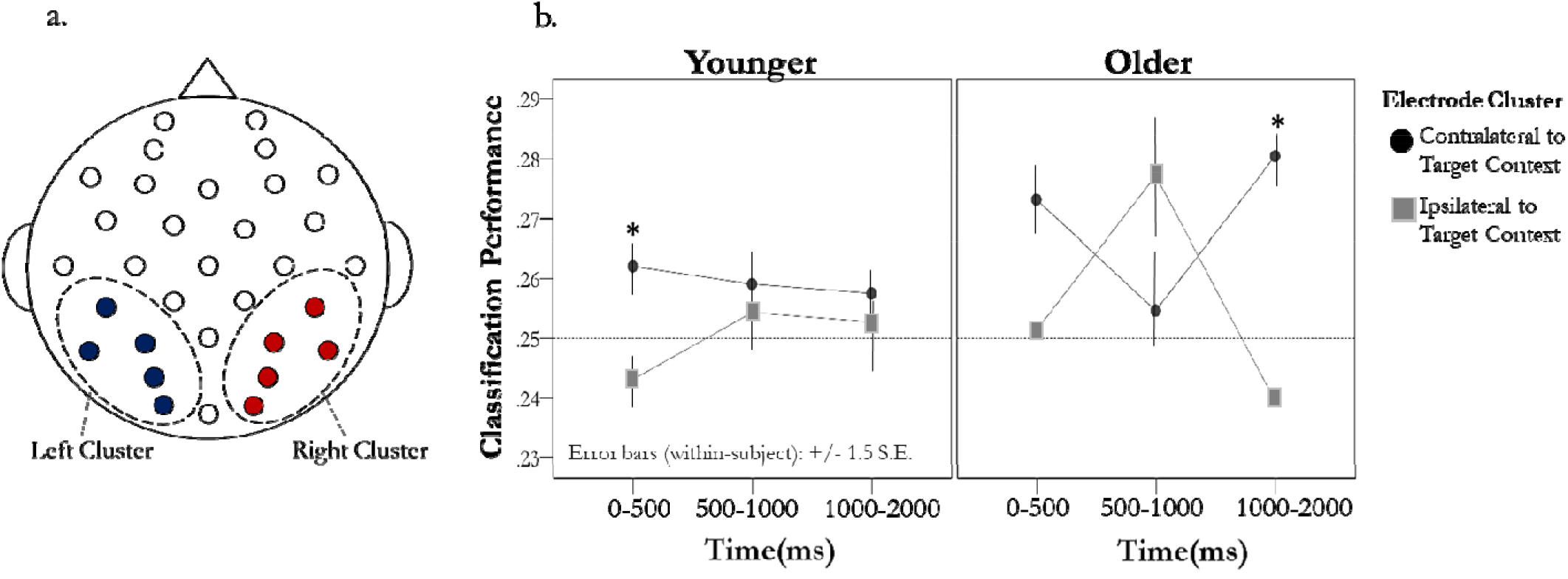
Electrode clusters and within-trial changes in classification performance. (a) Individual electrodes included in left and right posterior clusters. (b) Average classification performance across frequency bands at Time 1 (0-500ms), Time 2 (500-1000ms), Time 3 (1000-2000ms) in younger (left panel) and older adults (right panel). * indicates significant difference (p < .05) between contra- and ipsilateral cluster after correcting for multiple comparisons.

An initial repeated measures ANOVA, comparing Hemi-field (contralateral versus ipsilateral), Frequency Band (Delta, Theta, Alpha, Beta, Gamma), and Time (Time1, Time 2, Time 3) indicated a marginally significant main effect of hemi-field, *F*(1,39) = 3.19, *p* = .08, ƞ_p_^2^ = .08, but no main effect of frequency Band, *F*(1,156) = 0.60, *p* = .66, ƞ_p_^2^ = .02. In light of these findings, classification performance was averaged across frequency band prior to examining changes in classification performance within a trial. Although classifier performance was based on data aggregated over all conditions, to confirm that performance reflected both the spatial location and the type of target context feature (color or scene) we examined the probability estimates produced by the classifier (i.e., classifier evidence, see Supplementary material). The pattern of neural activity from these clusters not only reflected the spatial location of the target context, but also the specific type of context feature (e.g., color or scene; Supplementary **Figure S1**). Following this examination, the focus of subsequent analyses was determining the variation in classification performance within these clusters over time and between age groups.

To determine whether the performance difference between the contralateral and ipsilateral clusters varied as a function of age group over time we conducted a 2 (age) x 2 (hemi-field) x 3 (time bin) repeated measures ANOVA with a polynomial contrast. Results from this analysis indicated a significant hemi-field by time interaction, *F*(2,76) = 3.73, *p* = .03, ƞ_p_^2^ = .09. No other significant main effects or interactions were observed (*F*s ≥ 2.28, *ps* ≥ .12). To further examine the Hemi-field x Time interaction, we first compared classification performance in contralateral and ipsilateral clusters at each time point against chance (25%). After correcting for multiple comparisons, results indicated classification performance was significantly greater than chance in contralateral cluster at Time 1 [*M* = .27, *SD* = .05, *t*(39) = 2.63, *p* = .04] and marginally significant at Time 3 [*M* = .27, *SD* = .04, *t*(39) = 2.30, *p* = .06], whereas classification performance in the ipsilateral cluster was marginally significant at Time 2 [*M* = .27, *SD* = .05, *t*(39) = 2.19, *p* = .06]. While these findings suggest that electrodes contralateral to the target context’s spatial location may contain more category-specific information relative to ipsilateral electrodes, classification performance within these clusters was differentially affected over time and may reflect fluctuations in category-specific information for the target context.

### 3.5. Within-trial changes in Classification Performance and Context Memory Accuracy

Since the classification algorithm was trained to identify the target category representation, the within-trial decrease observed in the contralateral cluster around 500 to 1000 ms could reflect a degradation of the target category representation or an amplification of the distractor category representation that occurs when shifting one’s attention away from the target and towards the distractor. As such, we focused the following analysis on electrodes contralateral to the target context.

To explore how the observed changes in classification performance within a trial relate to context memory, we computed a difference score by subtracting classification performance at Time 1 (0-500 ms) from classification performance at Time 2 (500 – 1000 ms). Thus, negative values indicated a decrease in classification performance after the first 500 ms post stimulus onset and positive values indicated an increase in performance. The same calculation was used to examine the change in classification at the final time window (Time 3: 1000-2000 ms) by subtracting classification performance at Time 2 from classification performance at Time 3. Because the following analyses were exploratory, corrections for multiple comparisons were not carried out.

As can be seen in **Figure 8a**, greater declines in classification performance from Time 1 to Time 2 in the contralateral cluster (negative values) were associated with poorer target context accuracy, r(38) = .36, *p* = .02, and marginally greater distractor context accuracy, r(38) = −.28, *p* = .09. Changes in classification performance from Time 2 to Time 3 revealed similar trends [target context accuracy, r(38) = .23, *p* = .15; distractor context accuracy, r(38) = −.35, *p* = .03] (**Figure 8b**). While it is important to note that these relationships were primarily driven by older adults (**Figures 8a-8b**), in general, these findings show that declines in classification performance within an encoding trial result in poorer target and greater distractor context accuracy.

**Figure 8.**
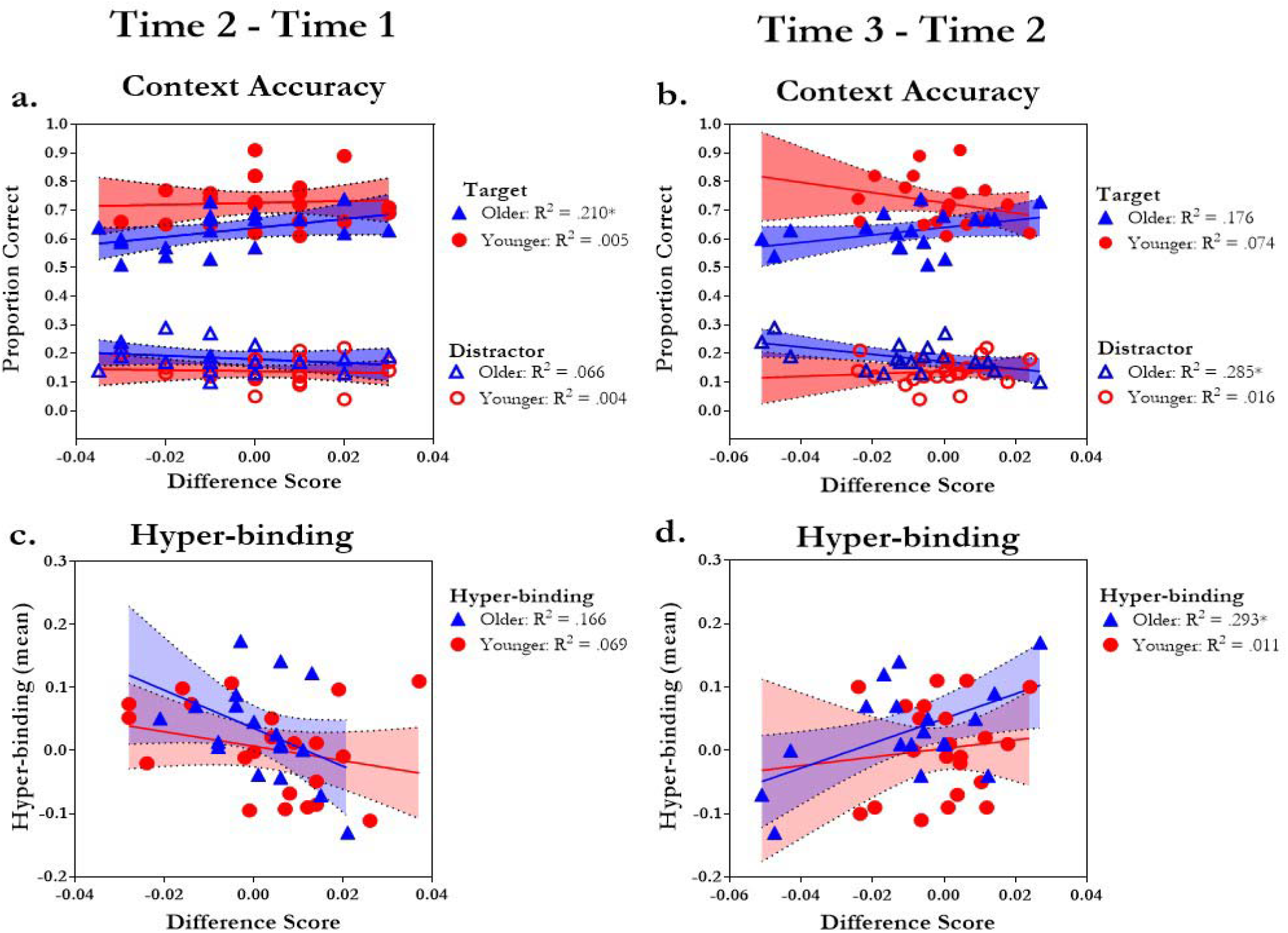
Within-trial changes in classification performance, context memory, and hyper-binding. Average target and distractor context accuracy (y-axis) with change in classification performance from (a) Time 1 (0-500ms) to Time 2 (500-1000ms) and (b) Time 2 (500-1000ms) to Time 3 (1000-2000ms). Average hyper-binding (y-axis) with change in classification performance from (c) Time 1 (0-500ms) to Time 2 (500-1000ms) and (d) Time 2 (500-1000ms) to Time 3 (1000-2000ms). *indicates significant R^2^ (*p* < .05).

One might assume that the pattern of reduced attention to the target category and/or amplification of the distractor would be predictive of greater hyper-binding. As shown in **Figure 8c**, greater declines in classification performance occurring approximately 0 to 500 ms after stimulus onset (i.e., Time 1 to Time 2) were associated with greater hyper-binding, r(38) = −.31, *p* = .05. In contrast, increases in classification performance from Time 2 to Time 3, were associated with more hyper-binding, r(38) = .28, *p* = .08, **(Figure 8d**). As can be seen in **Figures 8c and 8d**, these relationships were primarily driven by older adults.

## 4. Discussion

The goal of current study was to determine whether age-related differences in selective attention could explain patterns of context memory accuracy in older adults. In order to address issue, we applied multivariate pattern analyses (MVPA) on oscillatory EEG signals to track moment-to-moment shifts of attention between relevant and irrelevant contextual information during encoding. Findings provided evidence that patterns of neural oscillations reliably discriminated experimental conditions where attention was directed to a specific contextual feature (i.e., color or scene) in a specific spatial location (i.e., to the right or left of the center object) during encoding. Further, the degree to which neural oscillatory signals reliably differentiated between conditions within subjects predicted subsequent context memory accuracy across age. To our knowledge, this is the first report to use pattern classification techniques to decode selective attention in context memory encoding in younger and older adults, and provide evidence that individual and age-related differences in classification performance predict subsequent context memory accuracy and hyper-binding as measured during retrieval. The following sections will discuss the specific aspects of these findings, but first a brief review of the behavioral results is warranted.

### 4.1. Behavioral results

Younger and older adults exhibited similar item-memory performance. This permitted an exploration of age-related differences in context memory performance without the potentially confounding effect of age-related differences in item memory (Rugg and Morcom, 2005). For both groups, memory for contextual features was greater for the attended target context, relative to the unattended distractor context; older adults’ memory performance was near chance for the distractor context. Although these findings suggest both younger and older adults were successful at selectively attending to the target context during encoding, older adults exhibited poorer overall accuracy for the target context. Older adults also demonstrated numerically, although not statistically, higher hyper-binding of target and distractor context features, than young adults. We, and others, have previously shown statistically greater hyper-binding for older than younger adults (Campbell et al., 2012; Campbell et al., 2010; Campbell et al., 2014; James et al., 2016; Rowe et al., 2006). One possible explanation for the weaker effect observed in the current study is due to a slightly smaller sample of older adults in the present study relative to our previous report (see James et al., 2016). This, coupled with the fact that a small number of young adults show evidence of hyper-binding similar to many older adults, likely contributed to the small age group difference. Given these individual differences, we illustrate how individual differences in hyper-binding relate to individual differences in selective attention in the EEG oscillatory patterns below. However, we acknowledge that, on average, hyper-binding appeared to be more evident in older adults, and may be an indication of greater inhibitory deficits or difficulty ignoring or suppressing attention to the distractor context. That is, greater inhibitory deficits in older adults may lead to a reduced ability to encode the appropriate item-context relationship, resulting in poorer accuracy for the relevant context as well as greater hyper-binding (James et al., 2016; Strunk et al., 2017). As discussed below, the pattern classification results support this hypothesis.

### 4.2. Patterns of Alpha and Beta-band Oscillations Predict Context Memory Accuracy

From our previous analyses of ERPs and oscillatory signals at retrieval, we have suggested that poorer accuracy for relevant contextual information in older adults may be a function of greater inhibitory deficits during encoding. Here, we directly tested this hypothesis using classification analysis of EEG during encoding. In particular, classification performance within the beta band not only predicted the particular feature (scene or color) and location (left of right screen) of the target context, but was also strongly related to memory accuracy for the target and distractor context. Specifically, higher classification performance in the beta band was associated with greater target context accuracy and poorer distractor context accuracy, even after controlling for age. Similar relationships were also observed in the alpha band. Importantly, there was no relationship between individual differences in classifier performance and item recognition. This suggests that individual variability in classification performance is more likely related to the degree to which individuals are able to selectively attend to the target and ignore the distractor as opposed to general inattention, which would likely have a negative impact on item memory as well as item-context binding. Taken together, these results suggest that by taking the moment-to-moment variability in oscillatory EEG signals during encoding we can estimate the relative strength of target and distractor category representations that are within the focus of attention, representations which can then be used to predict subsequent context memory accuracy during retrieval. Furthermore, previous evidence has suggested that neural oscillations in Alpha and Beta bands serve different aspects of attention (Engel and Fries, 2010; Klimesch, 2012; Palva and Palva, 2007). For instance, Alpha-band oscillations have been associated with greater inhibition or suppression of attention to task-irrelevant information (Klimesch, 2012), whereas Beta-band oscillations have been associated with more controlled aspects of attention (Engel and Fries, 2010). Given the observed relationships between context memory accuracy and classification performance in Alpha and Beta bands, both inhibition of distractors and facilitation of targets may contribute to context memory accuracy in this study. However, the question as to which particular aspects of attention are captured in these frequency bands during context memory encoding will need further investigation.

### 4.3. Change in Classification Performance within and across Trials

Our examination of classification performance within a trial indicated changes in classification performance over time. Specifically, a decrease in classification performance in the electrode cluster contralateral to the target context was associated with poorer target context accuracy and greater distractor context accuracy. As previously mentioned, the classification algorithm was trained to identify the target context representation. As such, these within-trial decreases in classification performance may reflect degradation of the target context representation and/or amplification of the distractor context representation. Given that these relationships were primarily driven by older adults; it is possible that these changes in context representations were due to age-related differences in selective attention or inhibitory control.

It has been argued that reduced selective attention in older adults may be due to a broader attentional focus or “spotlight” compared to younger adults (Greenwood and Parasuraman, 1999, 2004). Specifically, some models suggest that an individual’s visuospatial attentional focus becomes broader with age, increasing the area of space that is scanned in an attempt to locate targets (Greenwood and Parasuraman, 1999, 2004). Consequently, older adults in the present study may have made more attentional shifts between target and distractor context features than did young adults. Importantly, although it is likely that aging may increase the number of saccades between target and distractor context features during encoding, we believe it is unlikely that the shift in classification performance across visual fields containing target and distractor contexts is solely the result of saccades for several reasons. First, we applied an ICA-based correction for saccadic movements prior to all time-frequency decomposition and classification analyses. Second, evidence suggests that the increase in one’s attentional focus with age can occur even in the absence of saccades (Greenwood and Parasuraman, 1999, 2004). Third, the posterior scalp distribution of the lateralized shift in classification performance is inconsistent with that of saccade-related EEG, which is more frontally-distributed (Huber-Huber et al., 2016). It is possible that the shift in classification performance could simply reflect a drift in attention away from the target, and not necessarily an explicit shift in attention toward the distractor. However, such an explanation is difficult to reconcile with the finding that this shift was associated with not only poorer target context accuracy but also greater distractor context accuracy, and greater hyper-binding.

Specifically, individuals that shifted their attention from away from the target approximately 500 to

1000 ms after stimulus onset (**Figure 6b**) were more likely bind distractors to targets (**Figure 7c**). Shifts that occurred between 1000 to 2000 ms were associated with less hyper-binding and greater accuracy for only the distractor context (**Figures 7b and 7d**). Since these fluctuations of attention to target features affected memory for both targets and distractors, we feel that the shift in classification performance may be better characterized as a shift in spatial attention from the target to the distractor context and then back to the target prior to individuals making their encoding judgments.

Although the poor spatial resolution of EEG does not allow us to assess the contribution of specific brain areas, we believe that inhibitory deficits are more likely to contribute to the present results for a few reasons. First, hippocampal binding impairments are thought to contribute to older adults’ miscombining features from similar, familiar events, which can manifest as highly confident false memories (Dodson et al., 2007a; Dodson et al., 2007b; Shing et al., 2009). In the current study, false alarm rates did not differ between young and older adults and confidence estimates were reduced, not increased, by age. Furthermore, while hippocampal binding deficits might contribute to poorer target context memory accuracy, it is not clear how they would lead to hyper-binding of relevant with irrelevant context features or fluctuations in attention to relevant context features within a trial during encoding.

Furthermore, while difficult to infer the underlying generators of scalp EEG signals, the posterior and lateralized distribution of this effect are consistent with parietal-occipital cortical generators sensitive to the locus of early visuospatial attention (e.g. Reynolds et al., 2000; Yantis, 2008). Imaging studies that have investigated the neural correlates of explicit and implicit memory have sometimes shown that early visual cortical processing may support implicit memory for object stimuli while additional frontal and medial temporal recruitment may be needed for explicit awareness (Koutstaal et al., 2001; Slotnick and Schacter, 2006). In addition to these previous findings, our behavioral results indicated that memory for the distractor context did not exceed the level of chance in older adults, fitting with the idea that hyper-bound associations between target and distractors may only be recognized implicitly. Taken together, we tentatively suggest that early visual processing of distracting context features is sufficient to bind them to simultaneously-presented targets but insufficient for them to be explicitly recognized.

Nevertheless, future neuroimaging studies that manipulate the level of between-event similarity and within event distraction will be informative for discriminating between these mechanisms underlying age-related source memory impairments.

Older adults’ reduced selective attention may also be affected by the mini-blocking method used in the current task to group trials according to the attended-to context. Aging may increase susceptibility to proactive interference from the previously relevant context feature during attempts to encode the currently relevant feature. Older adults may have trouble disengaging from the distractor category (i.e. scene or color) or a specific previously presented target feature (e.g. red) when it appears again as the to-be-ignored feature. Such an explanation would be consistent with evidence showing that older adults, to a greater extent than the young, form associations between pairs presented in close temporal proximity (Campbell et al., 2014) and with data showing that older adults’ working memory performance improves when proactive interference is minimized (Lustig et al., 2001). On the other hand, one might expect a build-up of proactive interference in older adults to result in overall poorer classification performance relative to younger adults. However, in the current study classification performance was equivalent in younger and older adults. Therefore, it may be best to consider proactive interference as potential contributing factor for poorer selective attention in older adults rather than a separate account. A future experiment in which trials were not blocked by target category type would allow for a direct test of the proactive interference hypothesis.

Thus far, we have proposed two attentional mechanisms (e.g., a broader attentional focus and/or proactive interference) that are consistent with the hypothesis that aging is associated with a diminished ability to suppress irrelevant information, which contributes to poorer episodic memory (Gazzaley et al., 2005; May et al., 1999). In some circumstances, this diminished capacity may result in the formation of excessive associations between context features. Although hyper-binding could conceivably benefit memory, such as in implicit memory tasks where retrieval of specific episodic details is not required (Campbell et al., 2010; Rowe et al., 2006), it is more likely to contribute to poor performance in traditional tests of memory and in real-world situations when these details are needed to distinguish one event from another (e.g., “Did I take my medication today or yesterday?”). However, previous evidence has suggested that selective attention in older adults can be improved with training (Mozolic et al., 2011). Therefore, additional training in selectively attending to relevant information and/or suppressing attention to distractors may further boost memory relevant contextual information and reduce binding of irrelevant contextual information.

In conclusion, the present study demonstrates how patterns of neural oscillatory signals can be used to decode selective attention to a specific context feature during a context memory encoding task and reliably predict subsequent memory of relevant context information during retrieval. During encoding, there was also evidence that selective attention fluctuated within a trial. These fluctuations were particularly evident in older adults and may reflect shifts of attention toward distracting information that impact memory for relevant context information. As such, these neural fluctuations may indicate hyper-binding. In sum, these results are consistent with emerging theories that age-related declines in context memory may be attributable to poorer selective attention and/or greater inhibitory deficits in older adults.

## Acknowledgements

This study was supported by National Science Foundation Grant # 1125683 awarded to Audrey Duarte. This publication was made possible with a Ruth L. Kirschstein National Research Service Award (NRSA) Institutional Research Training Grant from the National Institutes of Health (National Institute on Aging) Grant # 5T32AG000175. We thank all of our research participants

## Footnotes

1 To ensure the selection of 300ms time-bins did not unintentionally render more favorable classification performance, separate classification analyses were conducted with different time bins selected as features. Specifically, an across-electrode pattern using 16 time-bins (150ms per bin) and five frequency bands (e.g., Delta, Theta, Alpha, Beta, and Gamma) as features revealed a similar pattern of results to those reported in Figure 4b. That is, classification performance increased around 300ms and peaked approximately 600 to 1200ms. After correcting for multiple comparisons, this analysis indicated above chance (25%) classification performance at 600-750ms (*p*_corrected_ = .01) and 900-1050ms (*p* corrected = .02).

2 If task switching influenced the current pattern of results it is possible to have lower than expected classifier performance for the first few trials following a new task instruction (i.e., switching from judging the likelihood of color context feature to a scene context feature), particularly for older adults. Therefore, in a subsidiary analysis, we assessed classification performance over trials within each mini-block of task trials to determine if there were changes in performance over trials. Results from this analysis did not observe any significant differences between early and late trials within mini-blocks. Thus, in order to maximize power, we included all trials in the analyses.

3 The hyper-binding index was computed by subtracting the conditional probability of correctly endorsing the target context given the distractor was correct, p(Both correct)/[p(Both correct) + p(Distractor correct)], by the conditional probability of endorsing the target context given the distractor context was incorrect p(Target correct)/[p(Target correct) + p(Neither correct)]. Thus, positive values indicate a greater likelihood of correctly recognizing both the target and distractor (i.e., hyper-binding).

4 For consistency, we carried out a similar classification using the same eight time bins described in the previous across-electrode pattern (e.g., −300 to 0, 0 to 300, 300 to 600, 600 to 900, 900 to 1200, 1200 to 1500, and 1800 to 2000). The features included in this classification included two electrode clusters (left vs. right), eight time bins, and five frequency bands. The values for each feature represented average oscillatory power at each cluster-time-frequency band pairing. Classification performance and follow-up ANOVAs of this pattern yielded similar findings to those reported above.

